# Non-Canonical Cytochrome P450 Enzymes in Nature

**DOI:** 10.1101/2024.12.22.630014

**Authors:** Andy K. L. Nguy, Kendra A. Ireland, Chase M. Kayrouz, Juan Carlos Cáceres, Brandon L. Greene, Katherine M. Davis, Mohammad R. Seyedsayamdost

## Abstract

Cytochrome P450s (CYPs) are a superfamily of thiolate-ligated heme metalloenzymes principally responsible for the hydroxylation of unactivated C–H bonds. The lower-axial cysteine is an obligatory and universally conserved residue for the CYP enzyme class. Herein, we challenge this paradigm by systematically identifying non-canonical CYPs (ncCYPs) that do not harbor a cysteine ligand. Our bioinformatic search reveals 20 distinct ncCYP families with diverse ligands encoded in microbial genomes. We characterize a native serine-ligated CYP with a high-spin ferric resting state. Its crystal structure clearly shows a typical CYP fold and a serine alkoxide as a lower axial heme ligand. In addition, we report the discovery and characterization of the first native selenocysteine-ligated CYP in nature. Our findings radically expand the CYP metalloenzyme family.

## Main Text

Hemoproteins constitute a broad class of metalloenzymes that perform a diverse range of biological functions, including oxygen transport, electron transfer, and various essential enzymatic reactions in the cell^1-5^. The identity of the lower axial ligand to the heme dictates the function and reactivity of hemoproteins by modulating the electronic properties of the cofactor^6^. Several heme ligands are found in nature, including histidine in myoglobin (for oxygen binding)^7,8^, tyrosine in catalase (for peroxide disproportionation)^9,10^, and methionine in cytochrome c (involved in the electron transport chain)^1,12^. Among hemoproteins, cytochrome P450 enzymes (CYPs) form a large and distinct superfamily, members of which carry a specific protein architecture as well as an axial cysteine (Cys) ligand^13^. They catalyze myriad transformations, notably the remarkable stereospecific hydroxylation of chemically inert C–H bonds via oxygen rebound chemistry, as well as epoxidation, desaturation, and nitration reactions (**Fig. 1a, 1b**)^14-16^. CYPs are ubiquitous in all domains of life and play essential roles in cholesterol biosynthesis, antibiotic biogenesis, and drug metabolism^17-22^. All natural CYP homologs characterized to date feature a Cys-ligated heme, which governs P450 reactivity by providing the thermodynamic driving force, via the so-called ‘push-and-pull’ effect, to facilitate activation of strong C–H bonds^23,24^.

**Fig. 1.**
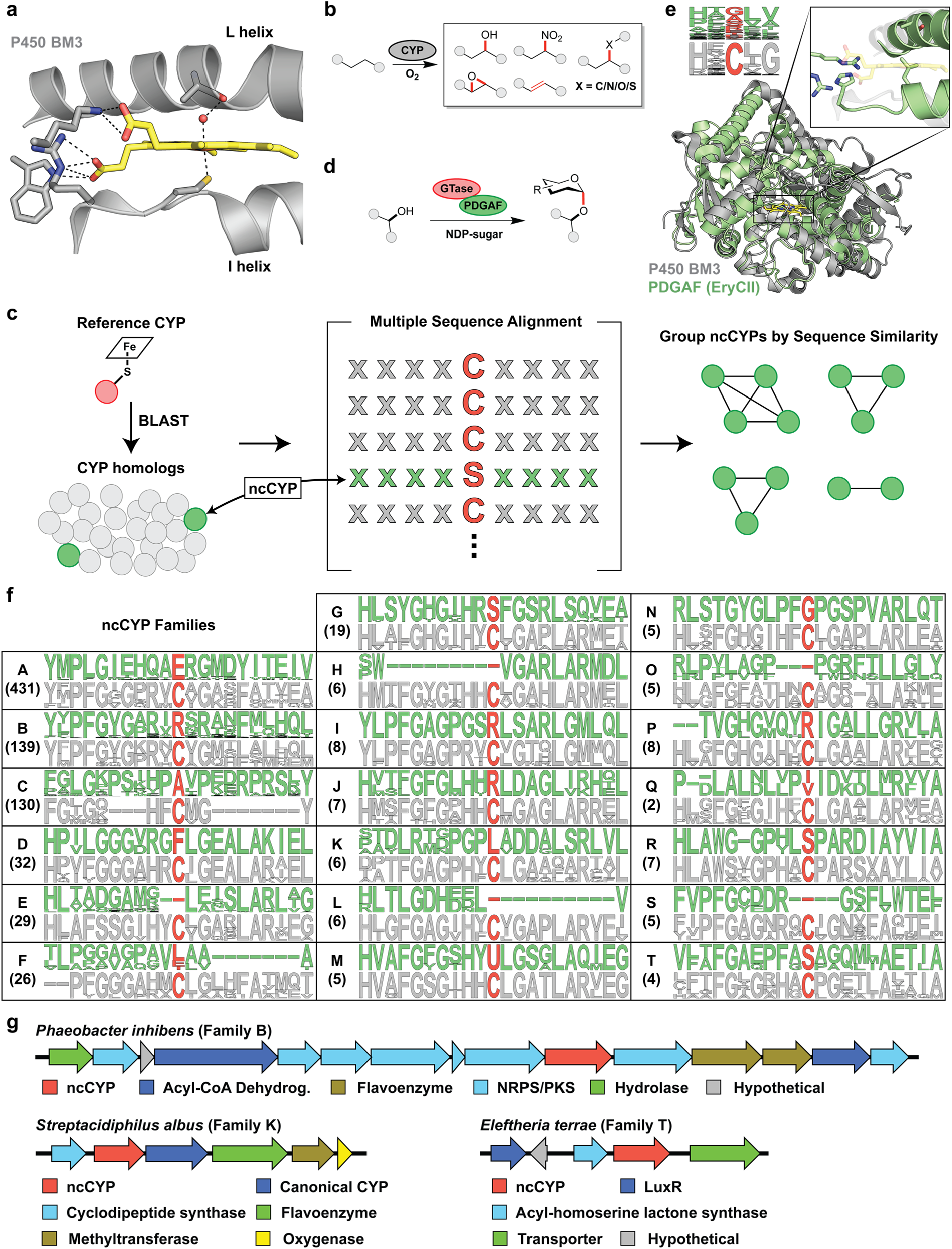
Bioinformatic analysis of non-canonical cytochrome P450 enzymes. **(a)** Representative CYP crystal structure (P450 BM3, PDB accession code 2IJ2) with lower-axial thiolate heme ligand. **(b)** Chemical transformations catalyzed by CYPs. **(c)** Bioinformatic strategy to identify ncCYPs through simple BLAST search, multiple sequence alignment, and sequence similarity analysis. **(d)** PDGAF acts as protein chaperone for glycosyltransferases. **(e)** PDGAF (EryCII, PDB accession code 2YJN) in green overlaid with heme-bound P450 BM3 in gray, and logo plots demonstrating sequence conservation surrounding the axial position (in red). **(f)** ncCYP families (in green), represented as logo plots illustrating sequence conservation surrounding the axial position, along with outgroups (in gray) representing the subfamily of canonical CYPs most closely related to each ncCYP family. Dashes represent alignment gaps. Family sizes are listed in parentheses. **(g)** Putative biosynthetic gene clusters that feature ncCYPs. Genes are color-coded as indicated.

Given the intimate relationship between the axial heme ligand and the properties of the cofactor, we wondered whether nature had selected for alternative ligands to modulate CYP function. To this end, we first employed a simple bioinformatic search strategy in which a small set of diverse CYP representatives was used as a query for BLAST alignment against the entire NCBI RefSeq sequence database (**Fig. 1c**). This resulted in over 200,000 BLAST hits, each of which represents a distinct CYP homolog from a sequenced organism. For each homolog, the residue at the putative axial position was inferred from its top scoring BLAST alignment. This approach relies on the assumption that the alignment position, but perhaps not the identity, of the axial ligand is invariant among all homologs. It almost certainly holds for the vast majority of CYPs, and we indeed find that >99.5% of all BLAST hits feature an expected Cys residue at the aligned axial position. Of the remaining sequences, 301 were removed due to their strong similarity to the PDGAF (P450-derived glycosyltransferase activating factor) family. These close homologs of CYPs carry the canonical protein fold but lack a heme cofactor and only serve as chaperones (**Fig. 1d, 1e**)^25,26^. Strikingly, upon removal of PDGAFs, we arrived at a list of 880 novel CYP homologs with an alternative residue at the aligned axial position. Genes encoding these unusual proteins are harbored by taxonomically diverse bacteria, spanning six phyla and over forty genera. The non-canonical CYPs (ncCYPs) were subsequently categorized into 20 subfamilies (designated as A through T) using sequence similarity analysis, revealing a rich assortment of putative axial ligands never before observed (**Fig. 1f**).

The two largest subfamilies (A and B) feature putative glutamate (Glu) and arginine (Arg) ligands, respectively, in place of Cys (**Fig. 1f**). Others exhibit serine (Ser, subfamilies G, R, and T) or selenocysteine (Sec, subfamily M) residues at the axial position, representing intriguing isosteric replacements of the canonical thiolate ligand. Residues with hydrophobic sidechains are found in several subfamilies as well, including alanine (subfamily C), phenylalanine (subfamily D), leucine (subfamilies F and K), and isoleucine/valine (subfamily Q). In some cases, no ligand is predicted (e.g. subfamilies E, H, L, O, and S). The fact that some residues appear in multiple clades suggests convergent evolution from disparate ancestors. Further analysis of the genetic context surrounding the ncCYPs revealed that several of the subfamilies are situated in operons containing other biosynthetic genes, suggesting their potential involvement in natural product biosynthetic pathways (**Fig. 1g**).

To ensure that these represent genuine CYPs with an alternative axial ligand, we first investigated a member from the putative Ser-ligated CYP family, a modification that would alter the electronic properties of the cofactor. Ser-ligated CYPs have been generated by site-directed mutagenesis from Cys-containing parental wild-type sequences, but they have never been characterized in native form. We recombinantly produced and purified the ncCYP from family G (termed P450-G9) encoded by *Actinokineospora terrae*. In close agreement with literature precedent on engineered Ser-ligated P450s by the Arnold group^27^, P450-G9 exhibits a strong Soret peak at 403 nm with a broad Q band at 605 nm and an additional feature at 485 nm (**Fig. 2a**). The dithionite-reduced ferrous P450-G9 shows a strong Soret peak at 423 nm. Upon formation of the ferrous-CO complex, this feature shifts to an intense band at 410 nm with well-defined Q bands at 534 nm and 567, in agreement with the engineered ‘P411’ enzyme^27^. Moreover, analysis by low-temperature X-band electron paramagnetic resonance (EPR) spectroscopy revealed an S = 5/2 high-spin ferric heme resting state^28^, with simulated broadband rhombic *g*-tensors of (7.0, 4.7 and 1.9) (**Fig. 2b**). These features are distinct from substrate-displaced high-spin CYP systems that have characteristic *g*-values near (8, 3.6, 1.7)^29^. A minimal amount of ferric low-spin population was also present for P450-G9, in stark contrast to the ferric low-spin values of (2.4, 2.2, 1.8) typical of substrate-free CYPs^30^. The high-spin ferric heme signature observed for P450-G9 is consistent with diminished sigma electron donation derived from the Ser ligand that enables a distorted 5-coordinate square pyramidal electronic geometry^28^. The spectroscopic data are thus consistent with the presence of a Ser-ligated heme.

**Fig. 2.**
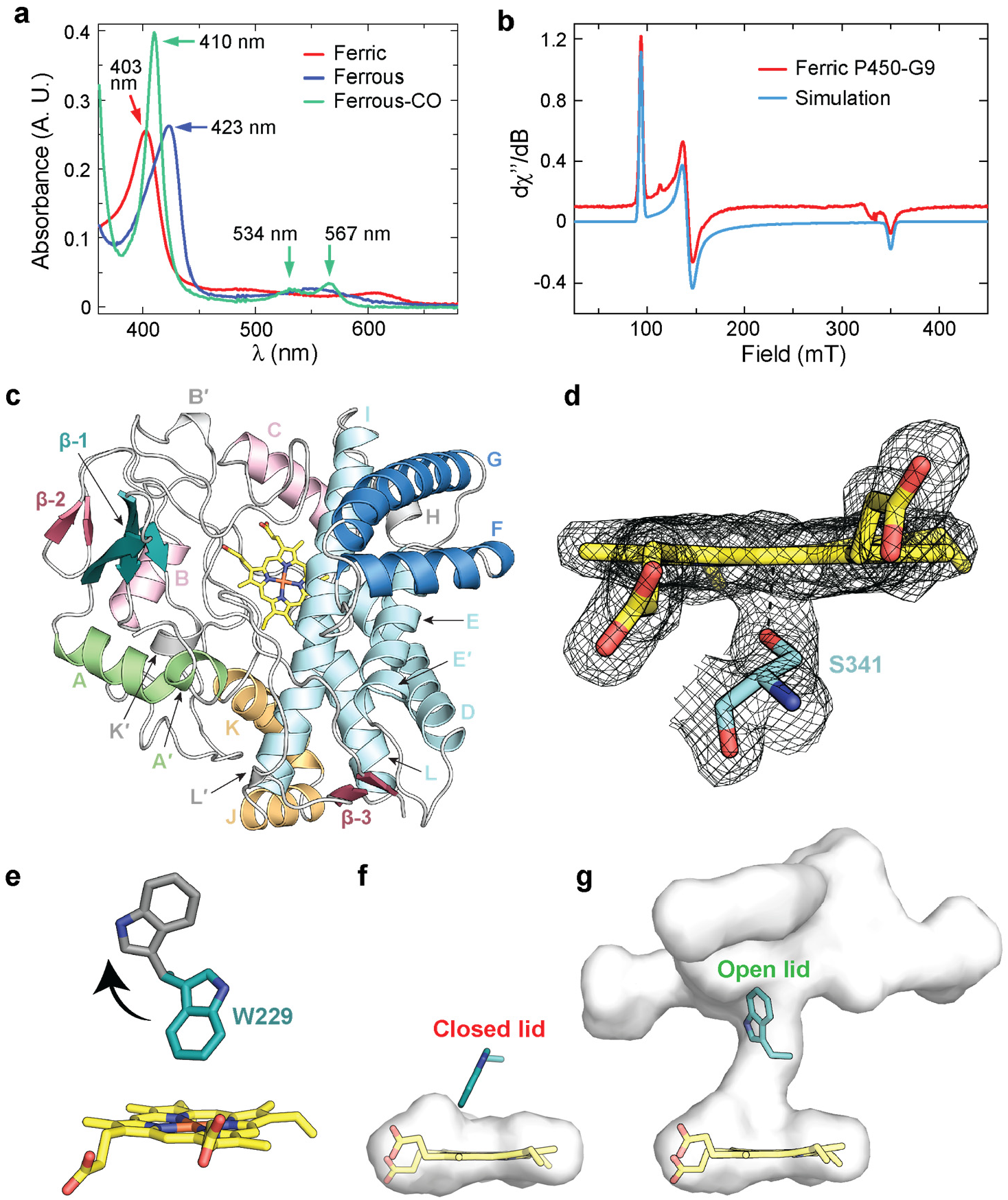
Spectroscopic and structural analysis of P450-G9, a Ser-ligated CYP. **(a)** UV/Vis spectra of P450-G9 in resting-state ferric form, reduced ferrous form, and ferrous-CO form. Key spectral features are marked. **(b)** X-band EPR spectrum and simulation of ferric high-spin P450-G9. **(c)** Structural overview of P450-G9 (PDB accession code 9CJG) with labeled secondary structures and the heme cofactor shown in yellow. **(d)** Alkoxide ligation to the heme cofactor via S341, as demonstrated by the 2Fo-Fc electron density map contoured at 1.0 sigma. **(e)** P450-G9 features a tryptophan residue in the center of the I helix, preceding a conserved alanine that typically hydrogen bonds to the water molecule occupying the sixth ligand position in cysteine-ligated CYPs. The crystallographically observed conformation of W229 (dark teal) blocks substrate binding and closes the distal solvent channel to heme. Rotation to the most common tryptophan rotamer (gray), however, would allow access to the iron center. **(f)** P450-G9’s solvent-inaccessible heme-binding pocket (523 Å^3^) with W229 in the observed closed-lid position. **(g)** Upon rotation of W229 to the open lid position, the heme-binding pocket becomes solvent accessible and increases nearly four-fold in volume (1965 Å^3^).

We subsequently solved the structure of P450-G9 via X-ray crystallography to further corroborate this conclusion (PDB accession code 9CJG). The resultant model, refined to 1.85 Å resolution, displays the hallmark CYP-fold (**Fig. 2c**), with clear electron density confirming the presence of a non-canonical alkoxide Ser ligand (residue S341) that forms a dative bond with the ferric heme center (**Fig. 2d**). Further inspection of the secondary coordination sphere reveals an unusual tryptophan residue (W229) lying near-perpendicular to the heme plane (**Fig. 2e**). Curiously, the indole side chain is positioned 4.7 Å from the heme iron and appears to block substrate binding. By adopting an unusual rotamer, it functions as an active site lid, sealing the heme-binding pocket from the external environment (**Fig. 2f**). Modeling of the most common W229 rotamer observed in protein structures (the so-called m95 rotamer) leads to an open active site, thus facilitating the creation of a narrow solvent channel to the heme cofactor (**Fig. 2g**). Volume calculations indicate that this putative side chain rotation results in a nearly four-fold increase in the active site volume (from 523 Å^3^ to 1965 Å^3^). Analysis of P450-G9’s binding pocket thus indicates that conformational changes may be imperative in enabling substrate ingress to the active site.

While uncommon, P450-G9 is not the only CYP that hosts a tryptophan residue distal to the heme cofactor. Prostacyclin synthase, a class III CYP that catalyzes the isomerization of prostaglandin H_2_, features a tryptophan at the same location in its I helix^31,32^. Unlike P450-G9, however, prostacyclin synthase’s analogous tryptophan does not block substrate binding, lying parallel with the heme plane and positioned 8.3–8.5 Å from the heme iron in substrate-free structures (PDB accession codes 2IAG and 3B98). Furthermore, the active site remains solvent accessible, and minimal changes to the protein structure are required for substrate binding. Collectively, these findings confirm identification of the first natural Ser-ligated CYP and implicate W229 and other structural features in regulating entry to the P450-G9 heme-binding pocket.

As a second case study, we further analyzed ncCYP subfamily M, putative selenoproteins featuring a Sec residue at the CYP axial position. Sec, often termed the 21^st^ amino acid^33^, is encoded by the opal stop codon (UGA) and ribosomally installed into proteins with the assistance of a specific elongation factor (SelB)^34^, which concurrently binds a structural mRNA element (selenocysteine insertion sequence, SECIS) downstream of the opal codon (**Fig. 3a**)^35^. There have only been three examples of artificially Sec-substituted CYPs reported by Ortiz de Montellano^36^, Hilvert^37^, and Green^38^, either through cysteine auxotrophy or an engineered SECIS-based strategy. To date, however, there are no known examples of a natural Sec-ligated CYP.

**Fig. 3.**
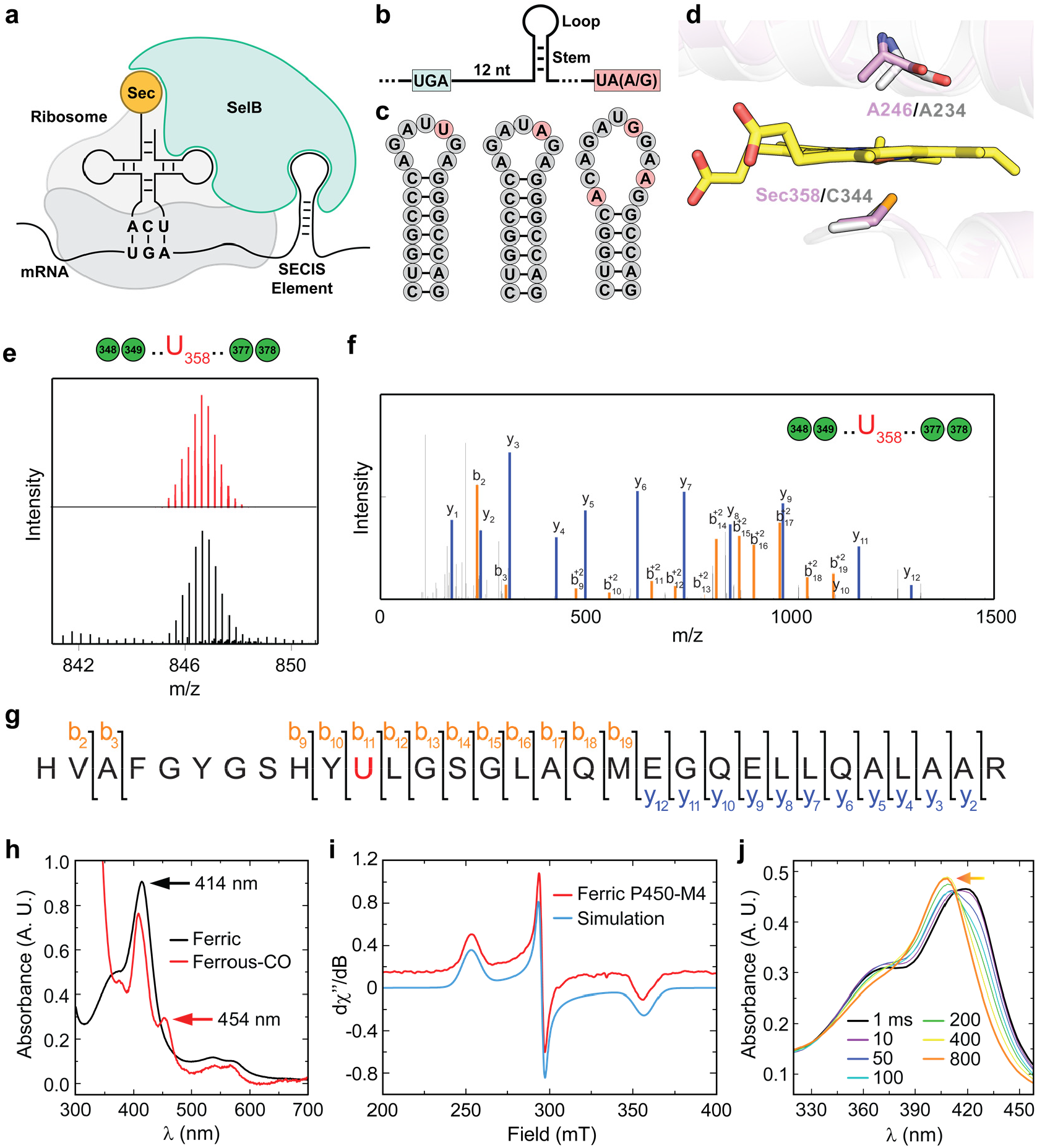
Selenoprotein biosynthesis and spectroscopic analysis of P450-M4, a Sec-ligated CYP. **(a)** Schematic representation of opal stop codon readthrough during selenoprotein biosynthesis, requiring the elongation factor SelB and downstream mRNA SECIS element. **(b)** Overall mRNA architecture of ncCYP family M transcripts, inferred from genome sequences. For each family member, the SECIS element is located 12 nt downstream of the opal stop codon (UGA). Open reading frames terminate with a non-opal stop codon (UAA or UAG). **(c)** Predicted mRNA SECIS elements within ncCYP family M transcripts exhibiting stem-loop topology. **(d)** Structural overlay of P450 107H1 (PDB 3EJD) with an AlphaFold2-generated model of P450-M4. The predicted Sec358 overlays with C344, a lower axial ligand in P450 107H1. **(e)** Observed (black) and simulated (red) isotope envelopes for the P450-M4 tryptic fragment containing Sec. **(f)** HR-MS/MS spectra exhibiting b and y ions for the P450-M4 tryptic fragment. **(g)** Inferred MS/MS fragments for the P450-M4 tryptic product from the data in panel **f. (h)** UV/Vis spectra of P450-M4 in the resting-state ferric form and the ferrous-CO form. Only partial reduction of Sec-ligated P450-M4 is observed under our experimental conditions. Key spectral features are marked. **(i)** X-band EPR spectrum and simulation of resting state ferric low-spin P450-M4. **(j)** SF UV/Vis spectra after reaction of P450-M4 with *m*CPBA. The spectrum after 1 ms (black) and 800 ms (orange) are shown as thick lines. The transition from the resting state to the new species is marked by the orange arrow.

Intriguingly, ncCYP family M, specific to a small clade of *Entotheonella* (marine sponge endosymbionts)^39,40^, appears to consist of Sec-ligated CYPs, evidenced by the presence of a conserved opal codon at the axial position as well as the full set of Sec-incorporation genes elsewhere on the chromosomes. Further *in silico* analysis through RNAfold^41^, an RNA secondary structure prediction tool, uncovered downstream stem-loop regions consistent with SECIS mRNA regulatory elements (**Fig. 3b, 3c**). Moreover, structural prediction by AlphaFold2^42^ reveals an overall topology consistent with canonical CYPs, including a near perfect superimposition of Sec with the axial cysteine ligand (**Fig. 3d**). Though conclusive validation of native opal codon readthrough is challenging due to the uncultivability of *Entotheonella*, our bioinformatic analyses strongly suggest Sec incorporation in these systems.

To gain more insights into this interesting subfamily, we selected one of the members, termed P450-M4, for biochemical characterization. Given the complex biochemical machinery required for Sec installation^43^, standard recombinant expression in *E. coli* is not possible. The Söll group recently pioneered an amber suppressor-based strategy to site-specifically install Sec in recombinant proteins, thereby eliminating the necessity for SelB/SECIS-mediated extension^44^. This method relies upon an engineered allo-tRNA^Sec^ that proficiently engages with *E. coli* Ser aminoacyl-tRNA synthetase and the canonical elongation factor Tu. Co-expression of SelA and SelD converts serine-acylated allo-tRNA^Sec^ to Sec^45^. To our delight, we observed robust selenoprotein expression when P450-M4 with a requisite amber stop codon was simultaneously co-expressed with a plasmid encoding SelA, SelD and allo-tRNA. Selenium quantification by ICP-OES elemental analysis revealed a yield of 0.63 ± 0.07 Se/mol protein and HPLC/HR-MS analysis confirmed the expected high-resolution mass of a tryptic fragment containing the Sec residue, concomitant with its distinctive selenium isotope envelope (**Fig. 3e**). Additionally, high-resolution tandem MS/MS fragmentation unequivocally demonstrated the site-specific incorporation of Sec at the expected position within P450-M4 (**Fig. 3f, 3g**). A UV/Vis absorbance spectrum of resting state ferric P450-M4 revealed a Soret peak at 414 nm and, upon reduction and reaction with CO, a ferrous-CO band at 454 nm (**Fig. 3h**). Given the enhanced nucleophilicity of Sec vs Cys, P450-M4 proved difficult to reduce even with excess strong reductants, such as titanium citrate; therefore, a significant amount of ferric P450-M4 remained after reduction and reaction with CO. The X-band EPR spectrum of P450-M4 exhibits a ferric, low-spin rhombic signature with g-values at (2.65, 2.27, 1.87) (**Fig. 3i**). These are in contrast to the values reported by Ortiz Montellano (2.46, 2.27, 1.98)^36^ and Hilvert (2.47, 2.27, 2.00)^37^, perhaps due to alterations in the secondary coordination sphere of the native Sec-ligated P450 that likely affect the properties and EPR parameters of the cofactor. Finally, we monitored the reaction of P450-M4 with the oxidant *meta*-chloroperoxybenzoic acid (*m*CPBA) to assess whether the reactive compound-I intermediate can be detected. Diode array spectra show a transition with a clear isosbestic point from the ferric resting state to a new species upon reaction with *m*CPBA (**Fig. 3j**). The diagnostic ∼690 nm feature, indicative of compound-I, was not observed as part of this new species; its features are instead reminiscent of the ferryl compound-Il intermediate^46^. The discovery of this native Sec-ligated P450 will facilitate further downstream studies to interrogate cofactor properties, reaction scope and mechanism, as well as possible biocatalytic applications.

Herein, we have bioinformatically predicted a collection of ncCYP enzymes, which carry a non-Cys residue at the lower axial position. Biochemical characterization confirmed the natural occurrence of Ser- and Sec-ligated CYPs. Others remain to be studied. Our results open a new field of inquiry in understanding the evolution, structure, and perhaps most importantly, the function of these alternative CYPs. Together, our findings challenge the dogma that CYPs must always possess a conserved Cys thiolate ligand. Electronic perturbations of the heme cofactor, via substitution of Ser, Sec, and other residues, are likely to modulate reactivity through push-pull chemistry. However, the exact function of uncharged residues, which could abolish the protein’s ability to bind the heme cofactor, remains unclear. Our findings further suggest that the CYP fold is a privileged protein scaffold, for which the axial ligand has been diversified throughout evolution to tune reactivity and modulate enzyme function. Future directions include elucidating the small molecules associated with the biosynthetic gene clusters featuring ncCYPs as well as unveiling the roles of the ncCYPs in these pathways. These and other studies are now enabled by the expanded chemical landscape of CYPs with exotic P450 heme ligands.

## Materials & Methods

### Materials

Ampicillin, IPTG, PMSF, lysozyme, potassium chloride (KCl), protease inhibitor cocktail, Sephadex G-25, Se and Fe ICP-MS standard (TraceCERT), tris(2-carboxyethyl)phosphine (TCEP), sodium selenite (Na_2_SeO_3_) and L-arabinose were purchased from Millipore-Sigma. LB broth, Terrific Broth, LB agar, δ-aminolevulinic acid hydrochloride, rhamnose monohydrate, DNAse I, tris base, imidazole, glycerol, potassium phosphate monobasic, potassium phosphate dibasic, sodium hydroxide, and HisPur^TM^ Ni-NTA resin were purchased from Fisher Scientific. Chloramphenicol and kanamycin were purchased from Apex Bioresearch Products. 50 KDa MWCO Vivaspin 20 filtration units and Vivaspin 20 pressure heads were purchased from Sartorius. Nickel nitrilotriacetic acid (Ni-NTA) agarose resin was purchased from Prometheus Protein Biology Products. SOC medium was purchased from Thermo scientific. Sequencing grade modified trypsin was purchased from Promega Corporation. Hi-Trap desalting 5 mL columns, were purchased from Cytiva Life Sciences. Milli-Q water (>17 MΩ) was used for preparing all solutions. ME6 cells^1^ (containing a scarless quadruple deletion ΔselABC ΔfdhF from *E. coli* strain BW25113) and plasmid pSecUAG-Evol2 (Addgene #163148) were a gift from Dr. Söll.

### General Procedures

All UV–vis spectra were acquired on a Cary 60 UV–visible spectrophotometer (Agilent). Low resolution high-performance liquid chromatography-mass spectrometry (HPLC-MS) analysis was performed on an Agilent instrument consisting of a liquid autosampler, a 1260 Infinity Series HPLC system coupled to a diode array detector, and a 6120 Series Single Quadrupole ESI mass spectrometer. High-resolution (HR) HPLC-MS was carried out on an Agilent UHD Accurate Mass quadrupole time-of-flight (Q-TOF) LC-MS system, equipped with a 1260 Infinity Series HPLC, an automated liquid sampler, a photodiode array detector, a JetStream ESI source and a 6540 Series UHD Accurate-Mass Q-TOF mass spectrometer. HPLC purifications were carried out on an Agilent 1260 Infinity Series analytical HPLC system equipped with a temperature-controlled column compartment, a diode array detector, an automated fraction collector, and an automated liquid sampler.

### Bioinformatics

All CYP homologs were retrieved from the NCBI RefSeq database through BLAST searches using a manually curated list of 50 diverse CYP representatives as query sequences. This resulted in a preliminary list of 230,189 BLAST hits. For each hit, the BLAST alignment was analyzed to determine the residue located at the axial position, resulting in 5,070 sequences with a putative non-cysteine residue at this position. The new list was then manually inspected to remove canonical CYPs resulting from a poor initial alignment (1,435 removed), bad sequences (i.e. truncations, obvious frameshifts, ambiguities, etc. [2,905 removed]), sequences with completely divergent/unalignable axial regions (140 removed), and sequences with high homology to the PDGAF family (301 removed). The remaining hits were then re-BLASTed against the full NCBI non-redundant database to obtain the final list of 880 ncCYPs, grouped by sequence similarity into 20 ncCYP families..

For each family, six canonical CYPs with the highest sequence similarity to its members were retrieved and defined as the outgroup. Each family and its associated outgroup was then subjected to multiple sequence alignment using MAFFT with the L-ins-I method, and the aligned regions surrounding the axial position (+/-10 residues) were visualized using the logomaker python script. All-by-all BLAST analysis for each family and outgroup was also performed, and results were visualized using Cytoscape. Genetic context for representative ncCYPs (15 kb up- and downstream) were retrieved using the NCBI E-direct toolkit. AlphaFold models for representative ncCYPs were used as input to the Dali server, and the top scoring structural homologs were overlaid and visualized in PyMol.

NcCYP family M, featuring an axial selenocysteine residue, was identified by the presence of a TGA stop codon in the axial position, represented as a truncated CYP sequence in the NCBI database. To obtain the full-length sequences, each was translated through the TGA codon until reaching the next stop codon, found to be either TAA or TAG in each case, suggesting that this is the true termination codon. SECIS elements downstream of the TGA codon were visualized using the RNAfold web server.

### Expression of Serine-Ligated P450-G9

Serine-ligated P450-G9 was expressed according to previously described procedures. Briefly, a 14-mL sterile culture tube containing 5 mL LB supplemented with Kan (50 µg/mL) was inoculated with a single colony of E. coli BL21(DE3) harboring G9-pET-28b. The 5 mL cultures were grown at 37°C and 200 RPM for at least 12 hours, at which point 500 µL (1% v/v) were used to inoculate 50 mL LB-Kan in a 250 mL Erlenmeyer flask. The intermediate culture was grown overnight at 37°C and 200 RPM, and subsequently used to inoculate large cultures. Erlenmeyer flasks (8 x 4L) 800 mL of TB supplemented with kanamycin were inoculated with 4 mL (0.5% v/v) of intermediate culture. The large-scale cultures were grown at 37°C/200 RPM to an OD600 nm ∼0.6, cooled in an ice bath for 5–10 minutes and supplemented with δ-aminolevulinic acid to a final concentration of 0.5 mM. The cultures were grown for an additional 20 min at 18°C/200 RPM, after which cultures were induced 0.2 mM IPTG. After 24 h, the cells were harvested by centrifugation (8000 x g, 20 minutes, 4°C), frozen and stored at -80°C. A yield of 10 g of cell pellet per L culture was typical.

### P450-G9 Crystallization

Crystals of P450-G9 were grown using the sitting drop vapor diffusion method at room temperature. To obtain crystals, 8 mg mL^-1^ P450-G9 in 50 mM Tris, 150 mM NaCl, pH 8 was mixed 1:1 with a precipitant solution of 60% (v/v) tacsimate pH 7, 0.7% (v/v) 1-butanol. Crystals with a rod-like morphology appeared within 2-5 days. For crystal harvesting, the crystals were looped and briefly transferred into cryoprotectant, comprised of the precipitant with 30% (v/v) ethylene glycol, before flash freezing in liquid nitrogen.

### P450-G9 X-Ray Data Collection and Processing

Diffraction data were collected at beamline 21-ID-F of the Advanced Photon Source at Argonne National Laboratory using a Rayonix MX300 detector. Crystals were maintained at 100 K to minimize X-ray-induced damage while images were collected sequentially (Δφ = 1°) with an incident wavelength of 0.97872 Å. The data were subsequently indexed and integrated using iMosflm. The intensity data were merged, scaled, and converted to structure-factor amplitudes using *POINTLESS, AIMLESS*, and *CTRUNCATE* within the CCP4 suite. Molecular replacement (MR) was first attempted with cytochrome P450 CYP107X1 (PDB accession code 7WEX), which has 51% sequence identity with P450-G9. The MR solution, however, significantly improved using the AlphaFold2 predicted structure as the search model in *PHASER*, after removing all residues from the model with per-residue confidence scores (pLDDT) below 90%. We therefore proceeded with the structure solved using the AlphaFold2 model. Model building was conducted in Coot, the structure refined in Phenix, and model quality assessed using Molprobity. The P450-G9 structure is in the orthorhombic P2_1_2_1_2_1_ space group and contains two molecules in the asymmetric unit. The final model (PDB accession code 9CJG) was refined to 1.85 Å resolution. Selected data processing and refinement statistics can be found in Table S3. Figures depicting the structure were generated with PyMOL. Surface representations and volumes of the heme-binding pockets were calculated using CavitOmiX (v. 1.0, 2022, Innophore GmbH).

### EPR Spectroscopy

Electron paramagnetic resonance (EPR) spectra were acquired at the Princeton University Department of Chemistry facilities. CW X-band EPR spectra were recorded at 10 K on a Bruker EMXplus EPR spectrometer equipped with an Oxford Liquid Helium cryostat. In general, the following parameters were used: power, 50 µW–10 mW; modulation amplitude, 5 G; modulation frequency, 100 kHz; center field, 3200 G; sweep width, 1500 G; sweep time, 60 s; 10 scans. All EPR spectra were simulated using the “pepper” utility from the EasySpin software package with the Kazan Viewer user-interface from the Alexey Silakov Lab in PennState.

### P450-M4 Amber Suppressor Reconstitution and Purification

*E. coli* ME6 were co-transformed with pSecEvol2 (with a chloramphenicol resistance marker) and pET28b-ch-p450-m4 by electroporation, cells were recuperated in SOC media for 2 hours at 37 °C and were inoculated into a 100 mL preculture of LB broth supplemented with 50 μg/mL of chloramphenicol and 50 μg/mL kanamycin. The preculture was grown overnight at 37°C with shaking at 200 r.p.m. The following day, 10 mL of the overnight preculture were inoculated into 4 × 1.5 L cultures of LB broth, supplemented with 35 μg/mL of chloramphenicol, 35 μg/mL of kanamycin, 15 μM of Na 2 SeO_3_ and 0.2% w/v L-Arabinose and grown at 37 °C, with shaking at 190 r.p.m. Protein expression was induced by adding 0.2 mM IPTG and 15 μM Na_2_SeO_3_ when the OD_600_ reached 1.0, then the culture was maintained at 25 °C for 48h. Cells were harvested by centrifugation at 8, 000 × g for 10 minutes, and the collected cell paste, approximately 3.5 grams wet cell paste per liter, was flash frozen in liquid N_2_ and stored at −80 °C until purification.

To protect selenocysteines from oxidative damage, selenoprotein purification was performed in an anaerobic vinyl glovebox (Coy Laboratory products Inc.) at <20 ppm O_2_. The frozen cell paste was brought into the glovebox and resuspended in resuspension buffer consisting of 50 mM potassium phosphate, 150 mM potassium chloride, 0.5 mM TCEP, 5% w/v glycerol, adjusted to pH 8.0. The resuspension was bought out of the glovebox and lysed in an Emulsiflex C3 homogenizer at 14,000 psi that was previously flushed with anaerobic buffer and maintained under Argon flow. Lysed cells were degassed in a Schleck line and brought into the glovebox and not exposed to oxygen thereafter. The cell extract was clarified by centrifugation in O-ring sealed bottles at 30,000× g for 1.5 hours and the cell debris was discarded. 20 mM Imidazole was added to the clarified lysate and then it was applied to a 20 mL Ni-NTA column pre-equilibrated with wash buffer composed of 20 mM imidazole, 50 mM potassium phosphate, 150 mM potassium chloride, 0.5 mM TCEP, 5% w/v glycerol, adjusted to pH 8.0. The resin was washed with 20 column volumes of wash buffer and eluted with elution buffer in which the imidazole concentration was raised to 200 mM. Orange colored fractions containing the protein were pooled and concentrated using 50 kDa MWCO Vivaspin 20 filtration units sealed with a Vivaspin 20 pressure heads to keep the concentrators oxygen free. Then the protein was buffer exchanged into resuspension buffer using a HiTrap desalting 5 mL column.

### Selenium Elemental Quantification

Selenium incorporation in purified selenoproteins was quantified by inductively coupled plasma optical emission spectroscopy (ICP-OES). We digested 9 nmol of protein by adding 86 μL of 70% trace metal free nitric acid and incubated overnight at room temperature followed by 2 hours incubation at 90 ºC. Once samples were cold, 70 μL of hydrogen peroxide was supplemented and incubated at 90 ºC for 1 hour. Water was then added to a final weight of 3 g total sample and analyzed in an Agilent 5800 ICP-OES in axial mode. Se in protein sample was determined by comparing to a calibration curve using Selenium standards (TraceCERT, Sigma-Aldrich) produced by dilutions in 2% nitric acid.

### Selenocysteine LC-MS/MS Analysis

Correct selenocysteine incorporation was assessed by liquid chromatography tandem mass spectrometry (LC-MS/MS). 100 μg of protein were unfolded in 20 μL of 8 M urea in 100 mM ammonium bicarbonate, and 10 mM DTT was added and incubated for 1 hour at 25 °C to reduce cysteines and selenocysteines. Cysteines and selenocysteines were alkylated with 15 mM iodoacetamide for 60 minutes in the dark, and the reaction was quenched by adding 10 mM DTT and incubating for 10 minutes. Next, the digested samples were diluted to <2 M urea using 100 mM ammonium bicarbonate and supplemented with 0.16 μg of trypsin. Reactions were incubated at 37 °C overnight. Reactions were quenched with a final concentration of 1% formic acid.

The digested samples (5 μg at 1 μg/μL) were injected into a Shimadzu LCMS-9030 equipped with an LC-ESI-Q-TOF system. The UPLC stationary phase was a Shim-pack Arata C18, 2.2 μm, 150 mm x 2.0 mm and the mobile polar phase a solution of H_2_O with 0.1% formic acid (A) and the non-polar mobile phase acetonitrile with 0.1% v/v formic acid (B). Peptides were resolved in a three-phase gradient from 1-19 % B over 15 mins, from 19-31% B over 66 minutes and 31-45% B over 10 minutes with a flow rate of 0.2 mL/min at 60 °C. Eluent from the LC was injected directly into the ESI-qToF. The mass spectrometer interface settings were as follow: nebulizing gas flow: 2 L/min, heating gas flow: 10 L/min, interface temperature 300°C and a desolvatation temperature of 526°C. The interface voltage was set to 4.5 V and the DL temperature to 250°C. The spectrometer was set in data dependent acquisition (DDA) in positive polarity using an event time of 0.25 s. 5 DDA events between 400 and 1500 m/z with a threshold of 200 counts and charges between 1 and 6 were selected for MSMS using a Q1 transmission window of 1 m/z. MSMS collision energy was set to 35 ± 17 V and MS2 ions were detected between m/z 200–1500.

The LC-MS/MS data was transformed using proteowizard MSconvert software and peptide search was performed using MSFragger in “Default search” mode using a database consisting of protein sequence and the next variable modifications on Cysteines: +57.02146 (Iodoacetamide), -15.977156 (Ser mutation), +104.96591 (Sec mutation + Iodoacetamide) and -33.98772 (Dehydroalanine) and also +15.9949 for Methionine oxidation and +42.0106 for N-term acetylation. The misincorporation of serine during protein translation can be explained, at least in part by, the modestly rate-limiting conversion of Ser allo-tRNA^Sec^ into Sec.

High resolution MS/MS spectra of the Serine and Selenocysteine peptides was obtaining by re-running the samples in the LC-MS in the same conditions as above, but the spectrometer was set in MS/MS detection mode in positive polarity, detecting MS/MS spectra from 100 to 2000 m/z with an event time of 0.5 s. Peptides corresponding with the masses detected using DDA mode (816.4089 m/z for Serine peptide and 846.6481 m/z for Selenocysteine peptide) were fragmented using a collision energy of 40 ± 20 V.

## Acknowledgments

The authors thank the National Science Foundation (GRFP No. 1937971 to K.A.I.), the Eli Lilly-Edward C. Taylor Fellowship in Chemistry (to C.M.K.), the Anid/Subdirección De Capital Humano/Beca de Doctorado Becas Chile/7220044 (to J.C.C.), and the National Institutes of Health (grants R35 GM154993 to B.L.G., R35 GM147557 to K.M.D, and R35 GM152049 to M.R.S.) for financial support.

## Competing interests

The authors declare no competing financial interests.

**Correspondence and requests for materials** should be addressed to Mohammad R. Seyedsayamdost

